# Meiotic drive against chromosome fusions in butterfly hybrids

**DOI:** 10.1101/2024.02.02.578558

**Authors:** Jesper Boman, Christer Wiklund, Roger Vila, Niclas Backström

## Abstract

Species frequently differ in karyotype, but heterokaryotypic individuals may suffer from reduced fitness. Chromosomal rearrangements like fissions and fusions can thus serve as a mechanism for speciation between incipient lineages but their evolution poses a paradox. How does underdominant rearrangements evolve? One solution is the fixation of underdominant chromosomal rearrangements through genetic drift. However, this requires small and isolated populations. Fixation is more likely if a novel rearrangement is favored by a transmission bias, such as meiotic drive. Here, we investigate transmission ratio distortion in hybrids between two wood white (*Leptidea sinapis*) butterfly populations with extensive karyotype differences. Using data from two different crossing experiments, we uncover a transmission bias favoring the fused state at chromosome with unknown polarization in one experiment and a transmission bias favoring the unfused state of derived fusions in both experiments. The latter result support a scenario where chromosome fusions can fix in populations despite counteracting effects of meiotic drive. This means that meiotic drive not only can promote runaway chromosome number evolution and speciation, but also that this transmission bias can be a conservative force acting against karyotypic change and the evolution of reproductive isolation. Based on our results, we suggest a mechanistic model for why derived fusions may be opposed by meiotic drive and discuss factors contributing to karyotype evolution in Lepidoptera.

## Introduction

Major chromosomal rearrangements leading to karyotypic differences can be important for the evolution of reproductive isolation and maintenance of species integrity. The underlying assumption to this argument is that heterokaryotypic individuals should experience reduced fertility as a consequence of meiotic segregation problems. While underdominant hybrid karyotypes may constitute powerful barriers to gene flow between divergent lineages (King 1993; Deineri et al. 2003), the evolution of karyotypic change is paradoxical. How can a chromosomal rearrangement reach fixation in a population when the heterokaryotype is underdominant? Theoretical work has shown that fixation of underdominant chromosomal rearrangements can occur in isolated populations with small effective population size (*N*_*e*_) where allele frequency change predominantly is caused by genetic drift (Lande 1979; Walsh 1982; Gavrilets 2004). For this reason, the generality of chromosome evolution as a mechanism for speciation has been questioned (Futuyma and Mayer 1980; Templeton 1981; Nei et al. 1983). However, the probability of fixation of an underdominant rearrangement will increase if the rearranged chromosome structure is favored by a transmission bias (White 1968), such as meiotic drive. A novel rearrangement will predominantly be found in heterozygous state. This is the critical stage for an underdominant rearrangement, since once it reaches an allele frequency of 0.5 it will experience the same average selection as the ancestral arrangement. A transmission bias, such as meiotic drive, may favor either the novel or the ancestral variant in heterokaryotypes and affect the fixation probability of chromosomal rearrangements. Meiotic drive can therefore either oppose or mediate the evolution of chromosome number differences and reproductive isolation between species.

Previous studies suggest that meiotic drive could be a common evolutionary force (Smith 1976; Henikoff et al. 2001; Pardo-Manuel de Villena and Sapienza 2001; Burt and Trivers 2006; Kern et al. 2015; Wei et al. 2017; Stewart et al. 2019). An observation supporting this hypothesis is that the number of acrocentric chromosomes per genome has a bimodal distribution in mammals, where most species have either only acrocentric or metacentric chromosomes (Pardo-Manuel de Villena and Sapienza 2001). If karyotype structure had evolved neutrally, we would rather expect a unimodal distribution of acrocentric/metacentric chromosomes. It has previously been shown that both centric-fusions and -fissions can be favored by meiotic drive (Pardo-Manuel de Villena and Sapienza 2001; Chmátal et al. 2014). Opportunity for drive in female meiosis arises due to polar body formation, i.e. the production of primordial egg cells that never get fertilized. Chromosomes that are preferentially segregating to the mature egg cell rather than the polar bodies will be transmitted to the offspring with a higher probability and can therefore increase in frequency in a population. In monocentric taxa, the spindle fibers attach to the centromere during meiotic division and differences between homologous chromosomes in kinetochore size may cause meiotic drive (Akera et al. 2017). Here, chromosomal rearrangements may play a role since fused and unfused chromosomes may differ in centromeric DNA content and recruitment of kinetochore proteins, which can lead to meiotic drive (Wu et al. 2018). While such “centromere drive” can result in karyotypic change, selfish centromeres seem to occur rather frequently and not only in fission/fusion heterokaryotypes (Henikoff et al. 2001; Dudka and Lampson 2022). This conclusion rests on the observation that both centromere sequences and the interacting kinetochore proteins have evolved rapidly in many taxa, while their function have been conserved (Henikoff et al. 2001). The molecular mechanism of centromere drive during female meiosis in a few monocentric organisms have been characterized in some detail (Chmátal et al. 2014; Akera et al. 2017, 2019; Clark and Akera 2021; Dudka and Lampson 2022). In contrast, little is known about the potential for meiotic drive and the underlying molecular mechanisms in holokinetic organisms, where centromere activity is distributed across numerous locations across the chromosomes (holocentric) during meiosis (Bureš and Zedek 2014).

Butterflies and moths (Lepidoptera) have received a lot of attention in cytogenetic studies, partly due to the possibility of using the karyotype for species characterization (Lorković 1941; Lukhtanov and Dantchenko 2002; Lukhtanov et al. 2005; Descimon and Mallet 2009; Vila et al. 2010; Dincă et al. 2011). Lepidopterans have holokinetic chromosomes in mitosis and meiosis (Maeda 1939; Suomalainen et al. 1973; Turner and Sheppard 1975). Most lepidopteran species have a chromosome number close to n = 31, but substantial variation exists (Lorković 1941; Lukhtanov 2014; de Vos et al. 2020). Macroevolutionary studies have shown that chromosome number variation is positively associated with the rate of speciation in some specific butterfly genera that have extensive karyotype differences between species (de Vos et al. 2020; Augustijnen et al. 2023). However, it is still unclear if the interspecific difference in karyotype is a result of genetic drift, natural selection, or some other fixation bias, such as meiotic drive. A few butterfly genera show especially extensive chromosome number variation. The wood white butterfly (*Leptidea sinapis*) has the greatest intraspecific variation in chromosome number of all non-polyploid eukaryotes. *Leptidea sinapis* individuals in Catalonia (CAT) have 2n = 106-108, while Swedish (SWE) individuals of the same species have 2n = 57, 58 (Lukhtanov et al. 2011, 2018). Most of the interpopulation differences in karyotype spring from derived chromosome fissions and fusions in the CAT and SWE population, respectively (Höök et al. 2023) and there is a cline in chromosome number between these two extremes across Europe (Lukhtanov et al. 2011). In spite of the remarkable amount of rearrangements, hybrids between SWE and CAT are fertile and viable with hybrid breakdown of viability in F_2_ and later generations indicative of recessive hybrid incompatibilities (Lukhtanov et al. 2018; Boman et al. 2023). These characteristics make *L. sinapis* an excellent model system for investigating the underlying evolutionary processes leading to karyotypic divergence. Hybrids are often used to investigate meiotic drive since drive systems are expected to rapidly lead to fixation or suppression by counter-adaptations (Hurst 2019; Fishman and Mcintosh 2019). In hybrids, dormant meiotic drivers may be released from suppression and drivers that have been fixed in the parental lineages may become observable due to reformation of heterozygosity (Phadnis and Orr 2009; Fishman and Mcintosh 2019). In addition, hybrids between SWE and CAT *L. sinapis* will be heterozygous for a large set of fissions and fusions. This can increase the overall power to detect transmission distortion, which may have a small effect on a per-generation timescale.

Here we performed crosses between SWE and CAT *L. sinapis* and sequenced a large set of F_2_ offspring to assess potential transmission distortion (i.e. deviations from strict Mendelian segregation), to determine whether meiotic drive may be acting in this system. Our aims were to answer two main questions: i) Is there evidence for transmission distortion for chromosomes of a certain rearrangement type (e.g. fusion in the SWE lineage)? ii) Is potential transmission distortion mediating or counteracting chromosome number divergence between populations?

## Materials and methods

### Crossing experiments

We performed two crossing experiments between SWE and CAT *L. sinapis* (Figure 1). First, pure lines of each population were crossed to form F_1_ offspring. Two ♀SWE x ♂CAT and five ♀CAT x ♂SWE F_1_ families were established by crossing offspring of wild-caught individuals from each parental line. Here, only females from the ♀CAT x ♂SWE survived until the imago (adult) stage. The F_1_ offspring were used to establish both an intercross (F_1_ x F_1_, n = 8) and a backcross F_2_-generation (F_1_ female x male SWE, n = 2). For the intercross F_2_ individuals, we monitored individual survival to determine the genomic architecture of hybrid inviability (see Boman et al. (2023), for more details). Here, all offspring (n = 599) were sampled, i.e. both those that survived until adulthood and those that died at some stage during development. For the backcross families, we sampled all eggs that each female laid, three days after egg-laying (n = 32 and n = 35, per female).

**Figure 1.**
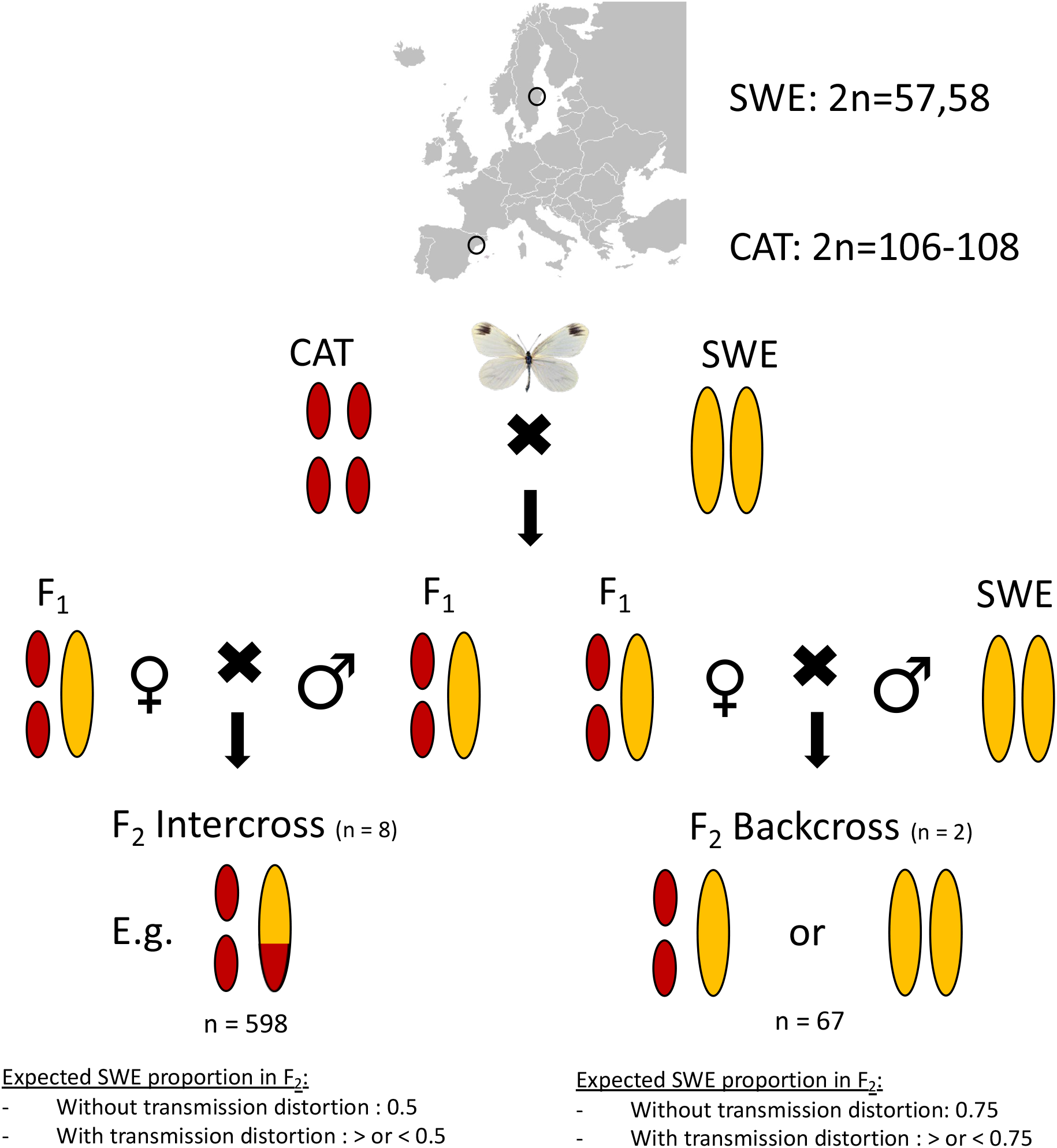
Crossing design and expected allele frequencies in the presence or absence of transmission distortion. Ovals represent an example of a homologous pair of autosomes. Note that female meiosis in butterflies is achiasmatic, i.e. recombination occurs only in males. Consequently, the F_2_ backcross is a test for female-specific transmission distortion.

### DNA extraction and pooled sequencing

We extracted DNA from the F_2_ hybrid offspring using a standard phenol-chloroform protocol. Individuals that died during development and eggs were extracted in pools of 2-21 individuals, due to low total DNA content in e.g. dead embryos. We measured the DNA content of each extracted sample using Qubit, and pooled samples to get equimolar concentrations of each respective individual. For the intercross, five different pools of F_2_ individuals were sequenced; dead embryos (n = 298), eggs (n = 73), dead larvae + dead pupae (n = 72), adult males (n = 76) and adult females (n = 80). The egg pool for the F_2_ intercross as well as eggs from the F_2_ backcross were sampled three days after laying. Pools were prepared for sequencing using the Illumina TruSeq PCR-free library preparation method and whole-genome re-sequenced (2 x 151 bp paired-end reads with 350 bp insert size) on a single Illumina NovaSeq6000 (S4 flowcell) lane at NGI, SciLifeLab, Stockholm.

### Inference of fixed differences

To measure transmission distortion in the offspring we used genetic markers and estimated allele frequency differences compared to the expected value based on each type of cross. We inferred fixed differences between the parental populations using population re-sequencing data from 10 SWE and 10 CAT male *L. sinapis* (Talla et al. 2019). In-depth information on variant calling can be found in Boman et al. (2023). Briefly, reads were trimmed and filtered and mapped to the Darwin Tree of Life reference genome assembly of a male *L. sinapis* from Asturias in north-west Spain, which is inferred to have a diploid chromosome number of 96 (Lohse et al. 2022). In total, we inferred 27,720 fixed differences and those were distributed across all chromosomes.

### Pool-seq read mapping and variant calling

We trimmed pool-seq reads and removed adapters using TrimGalore ver. 0.6.1, a wrapper for Cutadapt ver. 3.1 (Martin 2011). Seven base pairs (bp) were removed from the 3’ end of each read and all reads with an overall Phred score < 33 were discarded. Filtered reads were mapped to two modified versions of the reference genome assembly, where all fixed differences were set to either the SWE allele or the CAT allele, respectively. For subsequent analysis, we used the average allele frequency of both mappings to mitigate the effects of potential assembly biases. For the mapping, we used bwa *mem* ver. 0.7.17 (Li 2013). Mapped reads were deduplicated using Picard *MarkDuplicates* ver. 2.23.4 and reads with a mapping quality < 20 were discarded (Schlötterer et al. 2014). Variant calling was performed with MAPGD ver. 0.5 *pool* and only variants with a likelihood ratio score < 10^−6^ were retained (Lynch et al. 2014). In the presentation of the results, we arbitrarily decided to show the allele frequencies of the SWE allele for each respective marker in the pools of sequenced individuals. The number of loci that were retained for analysis after filtering were 27,713 in the backcross and 27,533 in the intercross experiment, respectively.

### Inference of transmission distortion

Rearrangement type classification was determined using parsimony based on synteny analyses between genome assemblies of *L. sinapis* and the related congenerics *L. reali* and *L. juvernica* (Höök et al. 2023; Näsvall et al. 2023). We inferred the degree of transmission distortion for four classes of rearrangements: derived fissions in the CAT population (Fission CAT), derived fusions in the SWE population (Fusion SWE), chromosomes with the two states segregating in all three *Leptidea* species (unknown polarization) and homologous autosomes. It should be noted that SWE has the fused and CAT the unfused state for all chromosomes with unknown polarization. We used these groups to increase the power for detecting small effect transmission distortions (see Table S2 for a list of sample sizes per group). Note that the *L. sinapis* karyotype includes three Z-chromosomes (Šíchová et al. 2015) and those were excluded since they are monomorphic for the SWE state in the backcross. To accommodate for the undefined order of events in complex rearrangements we restricted our analysis to chromosome units with a 1:2 ratio, i.e. where chromosome states in the two populations differ by a single fission/fusion event. Transmission distortion was evaluated using two-tailed binomial tests in *R* ver. 4.2.2 (R Core Team 2020). To produce counts of chromosomes from observed allele frequencies we rounded allele frequencies per pair for chromosomes with a fission/fusion rearrangement. Thus, for the sample size in the binomial tests, we counted pairs, since we conservatively assumed that the underlying mechanism (such as holokinetic drive) affects both unfused chromosomes equally and consequently there is only one event per homologous bivalent or trivalent during meiosis.

### Inference of ploidy

Patterns of transmission distortion can be caused by many processes, among them aneuploidy. We used pool-seq read counts at fixed differences to scan for the possibility of aneuploidy. If aneuploidy causes transmission distortion for a specific category of chromosomes, a higher sequencing read coverage for that category compared to other chromosome categories is expected. We therefore tested for significant differences in read coverage using both ANOVA and post-hoc analyses in *R*.

## Results

### Transmission distortion of derived fusions

We assessed potential transmission distortion in the F_2_ offspring from crosses between SWE and CAT *L. sinapis* using a pool-seq approach. The average allele frequencies in the F_2_ offspring for all marker loci (fixed alleles between the parental populations) were used to estimate potential deviations from strict Mendelian segregation using binomial tests. The analysis revealed significant transmission distortions for chromosomes with a derived fusion in the SWE lineage in both the F_2_ backcross (*p* ≈ 0.028) and the F_2_ intercross (*p* ≈ 0.024) (Table 1, Figure 1 and Table S3). In both cases, the unfused chromosome state characteristic for the CAT population was significantly overrepresented. This pattern was not driven by any specific outlier chromosome(s), since all except one chromosome (SWE) or chromosome pair (CAT) showed consistent deviations towards the CAT chromosome state (Figure 2). In the intercross, we also observed a significant transmission distortion for chromosomes with unknown polarization in the direction of the fused SWE state (*p* ≈ 0.003). Next, we considered explanations for the observed distortions. Since only Fusion SWE showed a significant deviation towards the CAT allele, it is not likely that the pattern is caused by reference bias. To test if aneuploidy could explain the observed transmission distortion, we calculated the coverage at marker loci for all chromosomes in the reference assembly (Figure S1). No significant differences between chromosome classes were observed, except between the Z chromosomes and the autosomes (Table S4), which is expected since the W chromosome is highly degenerated in Lepidoptera. This indicates that systematic aneuploidy is not causing the observed transmission distortion in our data.

**Table 1.** Expected and observed allele frequencies in the F_2_ backcross and intercross experiments and the results from binomial tests. Significant results are highlighted in bold.

| Experiment | Chromosome type | Expected frequency | Observed frequency | Lower 95% CI | Upper 95% CI | *p* value |
| --- | --- | --- | --- | --- | --- | --- |
| Backcross | Fission CAT | 0.75 | 0.761 | 0.712 | 0.806 | 0.659 |
| Backcross | Fusion SWE | 0.75 | 0.701 | 0.654 | 0.746 | **0.028** |
| Backcross | Homologous | 0.75 | 0.725 | 0.674 | 0.772 | 0.725 |
| Backcross | Unknown polarization | 0.75 | 0.731 | 0.674 | 0.783 | 0.481 |
| Intercross | Fission CAT | 0.5 | 0.497 | 0.479 | 0.516 | 0.798 |
| Intercross | Fusion SWE | 0.5 | 0.481 | 0.465 | 0.498 | **0.024** |
| Intercross | Homologous | 0.5 | 0.494 | 0.476 | 0.512 | 0.511 |
| Intercross | Unknown polarization | 0.5 | 0.531 | 0.511 | 0.551 | **0.003** |

**Figure 2.**
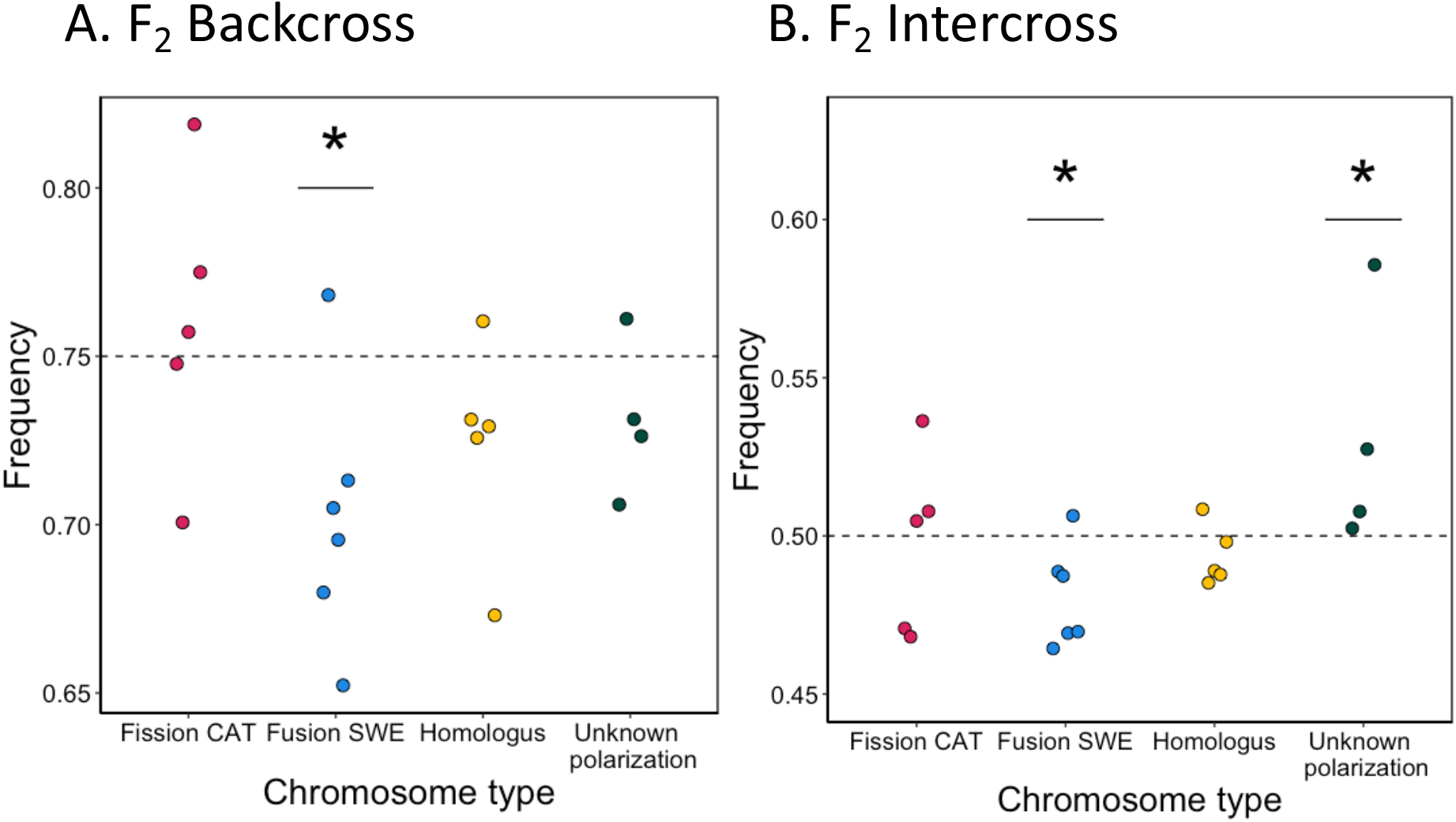
Average allele frequencies at marker loci for each chromosome (or pair of chromosomes for fission / fusion heterozygotes) in the F_2_ backcross (A) and the F_2_ intercross (B). In all cases, SWE has the fused state and CAT has the unfused state, except for the homologous (not rearranged) chromosomes, where both populations have the same state. Dashed lines represent the expected allele frequency in each experiment. Points have dodged positions along the x-axis to enhance visibility. Rearrangement types with significant transmission distortion are marked with an asterisk (*).

## Discussion

### Transmission distortion at derived fusions may be caused by female meiotic drive

Here we characterized transmission distortion using pool-seq of F_2_ offspring from crosses between SWE and CAT *L. sinapis*. We observed transmission bias in both crossing experiments at derived fusions, supporting the significance of the results. The fact that we observed a bias in the F_2_ backcross experiment suggest that female meiotic drive is causing the pattern at derived fusions. Mechanistically, the drive can be caused by differences in holokinetic binding of spindle fibers between the fused and unfused chromosome states, i.e. that the unfused ancestral state represented in the CAT population has stronger holokinetic activity. We only have rudimentary information available of the molecular components of the kinetochore structures and activities in Lepidoptera (Cortes-Silva et al. 2020; Senaratne et al. 2021). Like other holocentric insects, it seems that butterflies and moths lack the centromeric histone H3 variant (CenH3, also known as CENP-A), which is otherwise ubiquitous among eukaryotes (Drinnenberg et al. 2014). In mitotic cell lines from the silk moth, *Bombyx mori*, the kinetochore formation is directed towards heterochromatic regions of the chromosomes (Senaratne et al. 2021). If kinetochore activity is similarly associated with heterochromatic regions during female meiosis in F_1_ *L. sinapis* hybrids, it is possible that some unfused chromosomes have stronger centromeres due to proportionally more heterochromatin (Iwata-Otsubo et al. 2017). Chromosome fusion events might lead to loss of repetitive telomeric sequences at the fusion point (Figure 3A). In line with this, it has been shown that telomere-associated LINEs only constitute 5% of all LINEs close to fusion points in both *L. sinapis* and the congeneric *L. reali*, indicating that DNA has been lost in those regions (Höök et al. 2023). It should be noted that the genome assemblies used for that repeat analysis were based on 10X linked-read sequences and not long-reads. Since the assemblers using 10X linked-reads often fail to scaffold repeat-rich sequences (Peona et al. 2021), the amount of repetitive (and putatively heterochromatic) DNA at fusion breakpoints in *Leptidea* may therefore have been underestimated. If the meiotic drive observed for fused/unfused chromosome pairs is caused by differential kinetochore assembly due to loss of heterochromatin during fusion events, this can also explain why we did not detect any signal of meiotic drive for derived fissions. Fissions can form by double-strand breaks and are potentially not associated with the same heterochromatin differential between fused and unfused states. To conclusively test the hypothesis of holokinetic drive in *L. sinapis*, the next step will be to identify the kinetochore components and estimate the relative abundance of kinetochore proteins in meiotic cells in F_1_ hybrid females (Chmátal et al. 2014). Ideally, the kinetochore content can then be manipulated to experimentally validate if differential assembly of the kinetochore causes drive or not.

**Figure 3.**
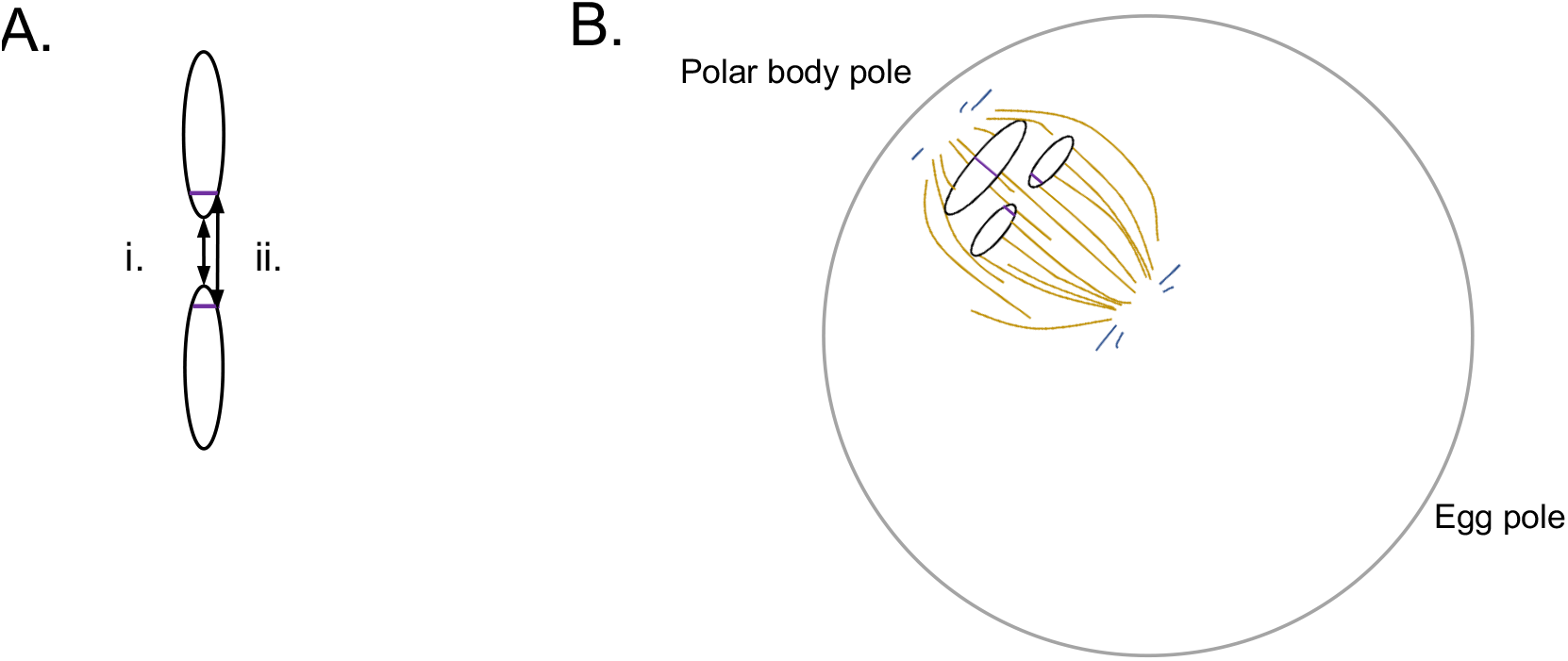
A model that describes how meiotic drive can occur during female achiasmatic meiosis of holokinetic organisms. (A) A fusion could either form through joining of ends (i) or e.g. non-homologous recombination, leading to loss of heterochromatic sequence at the fusion point (ii). (B) The loss of heterochromatic sequence could lead to a weaker holocentromere, which results in biased segregation during meiosis, either towards the polar body pole or the egg pole. If this mechanism explains the observed transmission distortion, the probability that the stronger holocentromere (in this case the unfused chromosomes) ends up in the mature oocyte is higher.

An alternative explanation to the observed transmission distortion would be early acting embryo viability selection enriched at chromosome fusions. While it is possible, we find it less likely since that would require that loci underlying viability are selected in both the F_2_ backcross and F_2_ intercross experiments, despite the different genomic backgrounds in individuals from those crosses. In addition, if two-locus hybrid incompatibilities cause such embryonic inviability in e.g. the F_2_ backcross experiment it would need to have a dominant gene action for the CAT allele (haplotype: CS/SS for the two loci), while at the same time have milder or no fitness consequences for the F_1_ parent with haplotype: CS/CS. While we cannot rule out such a scenario, we consider female meiotic drive to be a more parsimonious explanation for the biased allele frequency distributions observed here.

### Transmission distortion at segregating fission/fusion polymorphisms and the potential for male meiotic drive in Lepidoptera

We observed a transmission distortion favoring the fused state (SWE) for chromosomes with unknown polarization, i.e. rearrangement polymorphisms that are segregating within both *L. sinapis* and the closely related species *L. reali* and *L. juvernica*. This pattern is probably not caused by female meiotic drive since we did not observe such a transmission bias in the F_2_ backcross. This specific transmission distortion could potentially be caused by fertility selection on F_1_ parents which likely is stronger in F_1_ male than female meiosis in this system (Lukhtanov et al. 2018), early embryo viability selection, or drive during male meiosis. Lepidoptera have two distinct classes of sperm, nucleated (eupyrene) and anueclated (apyrene). Apyrene sperm have no nucleus and will therefore never contribute with genetic material to the next generation, similar to the situation for polar bodies in females (Friedländer 1997). This provides the opportunity for meiotic drive also in males, if for example specific chromosome arrangements have a higher probability to end up in eupyrene sperm. There could exist cheating mechanisms to avoid commitment from eupyrene to apyrene spermatogenesis, leading to meiotic drive among a heterozygous population of sperm.

### Causes and consequences of karyotype evolution in Lepidoptera

The potential for meiotic drive to cause karyotype evolution has been appreciated in both monocentric (Pardo-Manuel de Villena and Sapienza 2001) and holocentric organisms (Bureš and Zedek 2014). Here, we used a data set of 2,500 lepidopteran taxa (de Vos et al. 2020), to interpret our experimental evidence for transmission distortion for fission/fusion polymorphisms in *L. sinapis* (Figure 4). A visual inspection shows that a haploid count (n) of 31 chromosomes is the most common karyotype in Lepidoptera, but also that there is a substantial variation in chromosome numbers. Genera with species having a comparatively high number of chromosomes tend to have a higher variance in chromosome numbers (Figure 4, group i and ii). Only species within a few genera (*Leptidea* and *Polyommatus sensu lato*) have many members with high chromosome numbers (group i). A minority of species in group ii have n > 31 and a majority of genera comprise species with a maximum n <= 31 (group iii and iv). While no comprehensive phylogeny for the taxa included in this data set has been inferred, we can still use the information about chromosome number variation in Lepidoptera to draw a few conclusions. First, chromosome fusions are apparently widespread across Lepidoptera. This was recently confirmed by whole-genome alignments of more than 200 butterfly and moth species, (Wright et al. 2023). Recent models of chromosomal speciation and the role of chromosomal rearrangements in local adaptation have shown that a reduced recombination rate caused by a fusion event could be favored by selection and lead to speciation (Navarro and Barton 2003; Kirkpatrick and Barton 2006; Guerrero and Kirkpatrick 2014). Consequently, while meiotic drive could be involved it is not necessarily needed to explain the numerous chromosome fusions across the tree of Lepidoptera. Second, very few Lepidoptera species have high chromosome numbers as a consequence of multiple chromosome fissions. In both *Leptidea* and *Polyommatus*, which are the primary examples of species with highly fragmented karyotypes, inverted meiosis (i.e. sister chromatid segregation in meiosis I) has been observed (Lukhtanov et al. 2018, 2020a). It has been argued that while the achiasmatic (no crossover) female meiosis rescues fertility of trivalents, only holokinetic inverted meiosis rescues fertility (to some extent) in the chiasmatic male meiosis (Lukhtanov et al. 2018). Inverted meiosis in holokinetic organisms can thus reduce the selective disadvantage of trivalents in meiosis, increasing the probability for fixation of both fissions and fusions (Table 2). However, we do not yet know if inverted meiosis is a widespread phenomenon in Lepidoptera and how general such fertility rescue processes might be. In *Leptidea sinapis*, chromosome number is positively associated with the genetic map length (Näsvall et al. 2023), i.e. populations with more chromosomes have a higher recombination rate per physical unit length. An increased recombination rate as a consequence of chromosome fragmentation can potentially be beneficial, since a higher recombination rate reduces the impact of selection on linked sites (Fisher 1930). Signatures of linked selection has been documented in *L. sinapis* (Boman et al. 2021; Näsvall et al. 2023). However, an increased recombination rate also leads to a higher probability that beneficial associations between alleles in linked regions are broken up. We speculate that a higher chromosome number may also increase the risk of mis-segregation during meiosis. Given the potential costs of increasing chromosome number, it is possible that maladaptive meiotic drive has played a role in biasing the fixation of unfused chromosomes.

**Table 2.** Effects of different factors on karyotype evolution in Lepidoptera with special attention to the effects of meiotic drive.

| Factor | Effect | Consequence |
| --- | --- | --- |
| Epistatic selection | Selection for the co-inheritance of combinations of alleles on different chromosomes. | Decrease in chromosome number |
| Selective interference | Reduced efficacy of selection leading to selection for increased recombination | Increase in chromosome number |
| Holocentricity | Increased tolerance to chromosome fissions/fusions in female (achiasmatic) meiosis. | Increased variability in chromosome number. |
| Inverted meiosis | Rescued fitness of heterokaryotypes in male (chiasmatic) meiosis. | Increased variability in chromosome number. |
| Meiotic drive<br>(If supporting derived arrangement) | Fixation bias during female meiosis. | Increase or decrease in chromosome number. |
| Meiotic drive<br>(If supporting ancestral arrangement) | Fixation bias during female meiosis. | Conservation of chromosome number. |
| Meiotic errors | More chromosomes in meiosis leads to more opportunities for errors in meiosis. | Decrease in chromosome number |

**Figure 4.**
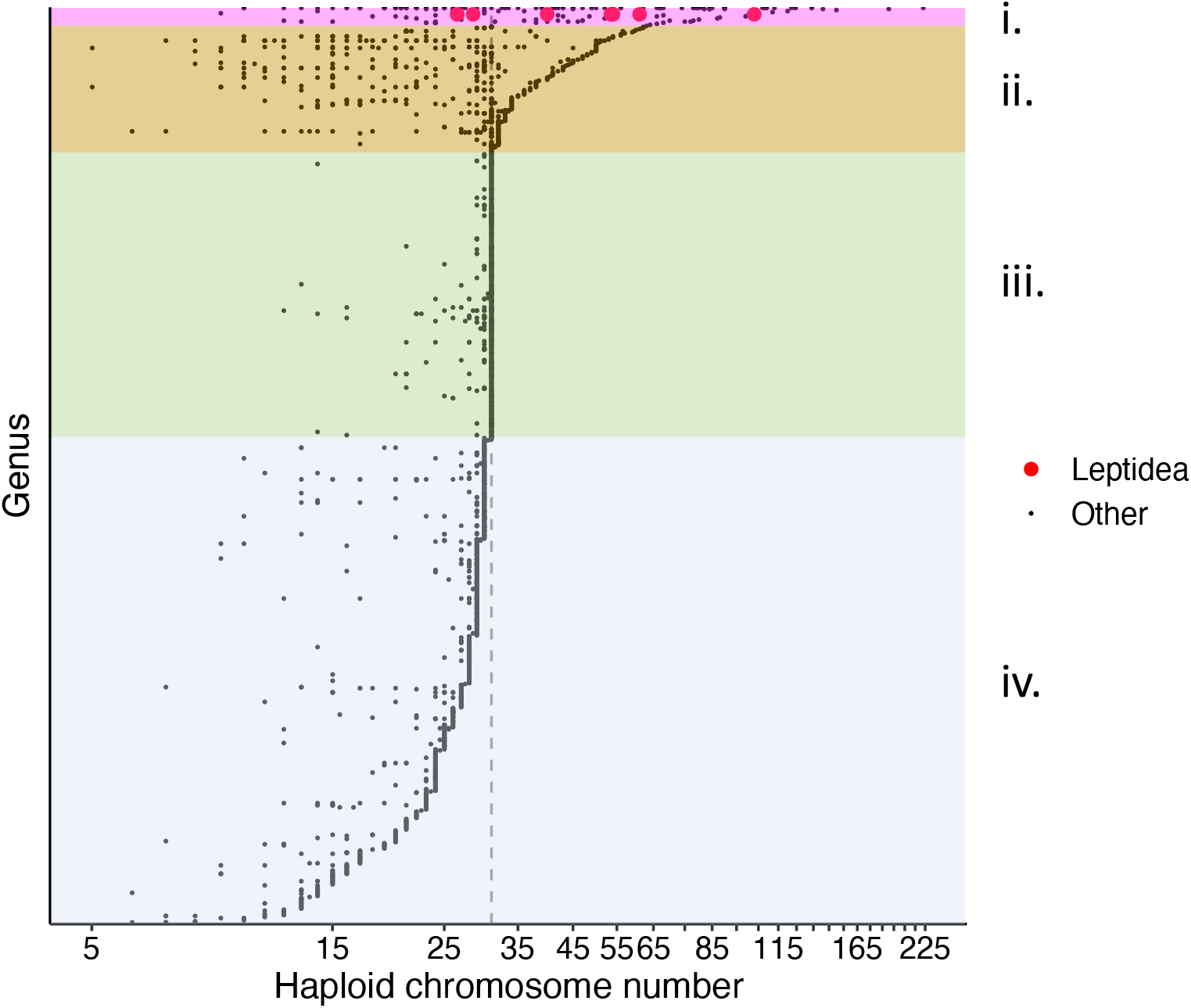
Haploid chromosome number count of 2,499 lepidopteran taxa from 869 genera. The data is from de Vos *et al*. (2020) with information from two *Leptidea* species added (Lukhtanov et al. 2011). The dashed vertical line indicates n = 31, the most common karyotype within Lepidoptera. Genera are sorted by maximum chromosome number with points representing individual taxa. Groups i-iv represents rough categories of chromosome number distribution per genus. Group i consists of a few genera with great within-genus variation in chromosome number and many members with n > 31. Group ii genera have high max counts and great within-genus variation, but the distribution is skewed towards low numbers. Group iii genera show generally low within-genus variation, and most members have n=31. Group iv genera have a max count < 31 with many genera having species with lower numbers.

### Meiotic drive opposing fixation of derived fusions

Since we observed a bias for the fused state for chromosomes with unknown polarization and the unfused state for derived fusions, predicting what continued intercrossing would do to chromosome number in this system is difficult. A tendency towards a higher chromosome number has been observed in crosses between Lepidoptera lineages with different karyotypes. In the closely related *Lysandra hispana* (n = 84) and *L. coridon* (n = 88 - 90), individuals tended to harbor the higher chromosome number after three generations of intercrossing (Beuret 1957). Similarly, in *Antheraea roylei* (n = 31) and *A. pernyi* (n = 49), intercrossed individuals in the F_23_ and F_32_ generations had n = 49 (Nagaraju and Jolly 1986). These results implicate that a fixation bias has been at play, since the expectation from genetic drift alone is the formation of a hybrid race with a karyotype distribution centered around the intermediate chromosome count (Lukhtanov et al. 2020b). In contrast to our study, the action of post-embryonic viability selection can however not be excluded in the crosses of *Lysandra* and *Antheraea*. In *L. sinapis*, we observed transmission distortion for derived fusions where the unfused chromosomes were overrepresented in the F_2_ offspring. This result does not support previously suggested models where meiotic drive promotes karyotype evolution (Pardo-Manuel de Villena and Sapienza 2001; Bureš and Zedek 2014). Instead, our results support a model where derived fusions are opposed by meiotic drive, i.e. that meiotic drive can act as a conservative force. If this pattern can be extrapolated more widely across Lepidoptera it lends further credence to positive selection acting on chromosome fusions, since they would have to fix while opposed by meiotic drive (Mackintosh et al. 2023). However, we emphasize that meiotic drive may very well have promoted karyotype change in some lepidopteran lineages (such as *Antheraea*), but conclusive experimental evidence for this is lacking. Experimental analyses across a wider range of taxa are needed to draw definitive conclusions on the general role of meiotic drive for karyotype evolution in Lepidoptera, but our results suggest that it may at least occasionally counteract karyotype change.

### Meiotic drive may be opposing evolution of hybrid inviability

In a previous study, we mapped the genomic architecture of F_2_ intercross hybrid inviability between the SWE and CAT chromosomal races of *L. sinapis* and observed a two-fold enrichment of candidate loci for hybrid inviability in derived fusion regions (Boman et al. 2023). This means that both transmission distortion and hybrid inviability are associated with the same chromosomes regions in this system, a pattern that has not been observed before as far as we know. However, genomic co-localization of regions affected by male meiotic drive and loci underlying hybrid sterility has been observed before in crosses between *Drosophila* taxa (Hauschteck-Jungen 1990; Tao et al. 2001; Phadnis and Orr 2009). It is believed that meiotic drive can promote the evolution of hybrid sterility through the formation of different driver-suppressor systems in divergent lineages experiencing limited gene flow (Frank 1991; Hurst and Pomiankowski 1991). Upon secondary contact, driver-suppressor systems could be misregulated and cause sterility in hybrids. While meiotic drive is intimately linked to reproductive processes, similar arguments could to some extent also be applied to hybrid inviability (Frank 1991; Hurst and Pomiankowski 1991). If meiotic drive accelerates sequence divergence, hybrid incompatibility could evolve as by-product of pleiotropy or physical linkage between the hybrid incompatibility locus and a driver or a suppressor. Conversely, since we observed meiotic drive in *L. sinapis* with a predisposition for the ancestral arrangement, it is possible that the factors contributing to hybrid inviability have evolved despite the counteracting force of meiotic drive. Consequently, the meiotic drive in the *L. sinapis* system could be opposing rather than promoting speciation. A similar pattern has previously been observed in *D. simulans* and *D. mauritiana*, where a driver has introgressed between species, which has resulted in reduced sequence divergence in that specific region (Meiklejohn et al. 2018). An alternative explanation would be that a substitution contributing to hybrid inviability reached high frequencies in the CAT population. Indeed, substitutions at Fusion SWE chromosomes in both populations could be contributing to hybrid inviability. More detailed characterization of the genetic basis of hybrid inviability is needed to further clarify the relationship between reproductive isolation and meiotic drive in this system.

## Data access

DNA-sequencing data is available at the European Nucleotide Archive under study id PRJEB69278. Scripts are available at GitHub in the following repository: https://github.com/JesperBoman/Transmission_distortion_Leptidea.

## Author contribution statement

JB and NB conceived and designed research. JB, NB and CW conducted experiments. JB analyzed data. JB wrote the manuscript with input from NB and RV. All authors read and approved the manuscript.

## Acknowledgements

This work was funded by Nilsson-Ehle Donationerna administered by the Royal Physiographic Society Lund (#146700346 to J.B.), the Swedish Research Council (VR research grant #019-04791 to N.B.) and by NBIS/SciLifeLab long-term bioinformatics support (WABI). R.V. was supported by grant PID2022-139689NB-I00 (funded by MCIN/AEI/10.13039/ 501100011033 and ERDF A way of making Europe), and by grant 2021 SGR 00420 (from Departament de Recerca i Universitats de la Generalitat de Catalunya). The authors acknowledge support from the National Genomics Infrastructure in Stockholm funded by Science for Life Laboratory, the Knut and Alice Wallenberg Foundation and the Swedish Research Council, and SNIC/Uppsala Multidisciplinary Center for Advanced Computational Science for assistance with massively parallel sequencing and access to the UPPMAX computational infrastructure.

