## Supplementary Information for "Meiotic drive against chromosome fusions in butterfly hybrids"

**Supplementary Figure 1**: (A) Coverage per chromosome in the F_2_ backcross experiment. (B) Coverage per chromosome in the F_2_ intercross experiment. Here, the weighted averages per chromosome across pools were used. Coverage was weighted by the sample size of each separate pool. In both experiments, no significant difference was observed among chromosome types, except that the Z chromosomes (Z_1_, and Z_2_) had as predicted lower coverage in some comparisons to other chromosome types.

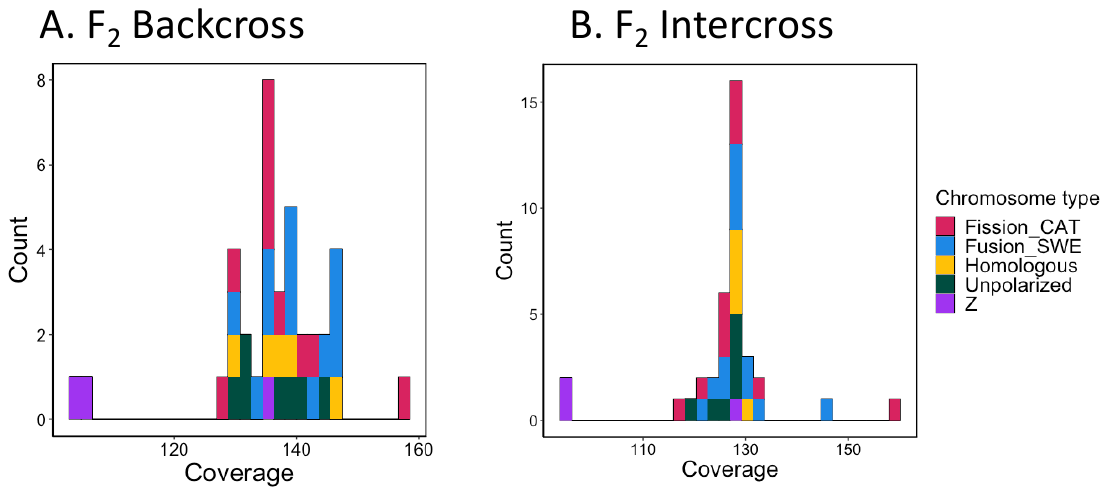

**Table S1**. Summary of sample sizes, sequencing yields and sequence quality scores for the F_2_ backcross pool-seq libraries. See Boman et al. (2023) for information on the F_2_ intercross pools.

| **Experiment** | **Pool** | **Sample size** | **Million reads** | **Percentage phred score**  **>=Q30(%)** | **Average**  **read depth with MQ>20*** |
| --- | --- | --- | --- | --- | --- |
| F_2_ backcross | Backcrossed eggs | 67 | 433.00 | 91.76 | 129x |

*At marker loci. MQ = mapping quality

**Table S2**. Sample sizes in the binomial tests used to quantify transmission distortion/meiotic drive.

| **Experiment** | **Chromosome category** | **Binomal test sample size** |
| --- | --- | --- |
| Backcross | Fusion SWE | 67*6 = 402 |
| Backcross | Fission CAT | 67*5 = 335 |
| Backcross | Unknown polarization | 67*4 = 268 |
| Backcross | Homologous | 67*5 = 335 |
| Intercross | Fusion SWE | 599*6 = 3594 |
| Intercross | Fission CAT | 599*5 = 2995 |
| Intercross | Unknown polarization | 599*4 = 2396 |
| Intercross | Homologous | 599*5 = 2995 |

**Table S3**. Estimated average allele frequency at marker loci per chromosome (or chromosomes) in the *L. sinapis* reference genome. Green shaded are chromosome(s) showing same direction of deviation from expected allele frequency in both experiments. The most notable difference was the shift towards the fused state in the F_2_ intercross for several of the “Unknown polarization” category. Differences between experiments could be due to either sampling error or e.g. biological differences stemming from the greater number of allelic combinations possible in the F_2_ intercross.

| **Chromosome(s)** | **Chromosome category** | **Allele freq. F_2_ backcross** | **Allele freq. F_2_ intercross** |
| --- | --- | --- | --- |
| 7+24 | Fusion SWE | 0.680 | 0.464 |
| 16+27 | Fusion SWE | 0.768 | 0.489 |
| 20+31 | Fusion SWE | 0.705 | 0.487 |
| 14+35 | Fusion SWE | 0.695 | 0.469 |
| 25+37 | Fusion SWE | 0.652 | 0.506 |
| 45+47 | Fusion SWE | 0.713 | 0.470 |
| 17+34 | Fission CAT | 0.748 | 0.471 |
| 40+15 | Fission CAT | 0.701 | 0.468 |
| 39+38 | Fission CAT | 0.757 | 0.505 |
| 10+18 | Fission CAT | 0.819 | 0.536 |
| 5+28 | Fission CAT | 0.775 | 0.508 |
| 33+8 | Unknown polarization | 0.706 | 0.502 |
| 41+46 | Unknown polarization | 0.761 | 0.508 |
| 4 | Unknown polarization | 0.731 | 0.527 |
| 44+36 | Unknown polarization | 0.726 | 0.586 |
| 23 | Homologus | 0.731 | 0.508 |
| 19 | Homologus | 0.726 | 0.485 |
| 11 | Homologus | 0.760 | 0.489 |
| 12 | Homologus | 0.729 | 0.488 |
| 1 | Homologus | 0.673 | 0.498 |

**Table S4**. Tukey honest significance difference tests for differences in average coverage between chromosome categories at marker loci. Significant p-values are highlighted with bold face font.

| **Experiment** | **Comparison** | **Difference** | **Lower** | **Upper** | **Adjusted *p*** |
| --- | --- | --- | --- | --- | --- |
| Backcross | Fusion_SWE-Fission_CAT | 1.94 | -7.48 | 11.36 | 0.97 |
| Backcross | Homologous-Fission_CAT | -0.56 | -12.61 | 11.48 | 1.00 |
| Backcross | Unknown_polarization-Fission_CAT | -1.72 | -12.56 | 9.12 | 0.99 |
| Backcross | Z-Fission_CAT | -22.91 | -37.39 | -8.43 | **0.00** |
| Backcross | Homologous-Fusion_SWE | 2.50 | -14.21 | 9.21 | 0.97 |
| Backcross | Unknown_polarization- Fusion_SWE | -3.66 | -14.12 | 6.80 | 0.85 |
| Backcross | Z-Fusion_SWE | -24.85 | -39.05 | -10.66 | **0.00** |
| Backcross | Unknown_polarization-Homologous | -1.16 | -14.04 | 11.72 | 1.00 |
| Backcross | Z-Homologous | -22.35 | -38.41 | -6.29 | **0.00** |
| Backcross | Z- Unknown_polarization | -21.19 | -36.37 | -6.02 | **0.00** |
| Intercross | Fusion_SWE-Fission_CAT | 0.22 | -10.29 | 10.74 | 1.00 |
| Intercross | Homologous-Fission_CAT | -0.55 | -14.00 | 12.91 | 1.00 |
| Intercross | Unknown_polarization-Fission_CAT | -3.62 | -15.72 | 8.49 | 0.91 |
| Intercross | Z-Fission_CAT | -23.09 | -39.26 | -6.92 | **0.00** |
| Intercross | Homologous-Fusion_SWE | -0.77 | -13.85 | 12.31 | 1.00 |
| Intercross | Unknown_polarization- Fusion_SWE | -3.84 | -15.53 | 7.84 | 0.87 |
| Intercross | Z-Fusion_SWE | -23.32 | -39.17 | -7.46 | **0.00** |
| Intercross | Unknown_polarization-Homologous | -3.07 | -17.46 | 11.31 | 0.97 |
| Intercross | Z-Homologous | -22.55 | -40.49 | -4.61 | **0.01** |
| Intercross | Z- Unknown_polarization | -19.48 | -36.43 | -2.52 | **0.02** |
